# The structure and function of the sucker systems of hill stream loaches

**DOI:** 10.1101/851592

**Authors:** Jay Willis, Theresa Burt de Perera, Cait Newport, Guillaume Poncelet, Craig J Sturrock, Adrian Thomas

## Abstract

Hill stream loaches (family *Balitoridae* and *Gastromyzontidae*) are thumb-sized fish that effortlessly exploit environments where flow rates are so high that potential competitors would be washed away. To cope with these extreme flow rates hill stream loaches have evolved adaptations to stick to the bottom, equivalent to the downforce generating wings and skirts of F1 racing cars, and scale architecture reminiscent of the drag-reducing riblets of Mako sharks. Hill stream loaches exhibit far more diverse flow-modifying morphological features than fast pelagic predators, suggesting as yet unknown drag reducing systems remain to be discovered. Here we describe the skeletal structure of *Sewellia lineolata* and *Gastromyzon punctulatus* and contrast that with other fish that face similar hydrodynamic challenges. We identify a major structural variation within *Balitoridae* pelvic sucker attachment positions which may explain fundamental constraints on the parallel development of different genera and which has not been described before. We also use high speed video capture, CT scans and Frustrated Total Internal Reflection to image and measure the sucker system in live operation and describe how it functions on a familiar activity for hill stream loaches (climbing waterfalls). We show how they can drag 3 to 4 times their own bodyweight up a vertical glass waterfall. Adaptations to high flow rates are the inspiration for this study, because there are many engineering applications where the ability to deal with high flow rates are important - either by reducing drag, or by generating the forces needed to hold an animal in place.

## Introduction

Hill stream, butterfly or sucker loaches (families, *Balitoridae* and *Gastromyzontidae*) are endemic to South Asia. In particular around the countries bordering the South China Sea (Kottelat, 2012). Often individual species are limited to a single catchment, or to several neighbouring catchments. The loaches found in neighbouring catchments often have identical morphology, although there are also broad similarities between more distant species. In general they have flattened ventral surfaces with body suckers and rasping mouths on their ventral surface (Kottelat, 2012). They also share an absence of fin spines, claws, adapted teeth (odontodes), or other claw like adaptations for station holding in currents which are common among other fishes and animals which inhabit similar mountain streams (Fig. 1) (De Meyer and Geerinckx 2014).

**Figure 1.**
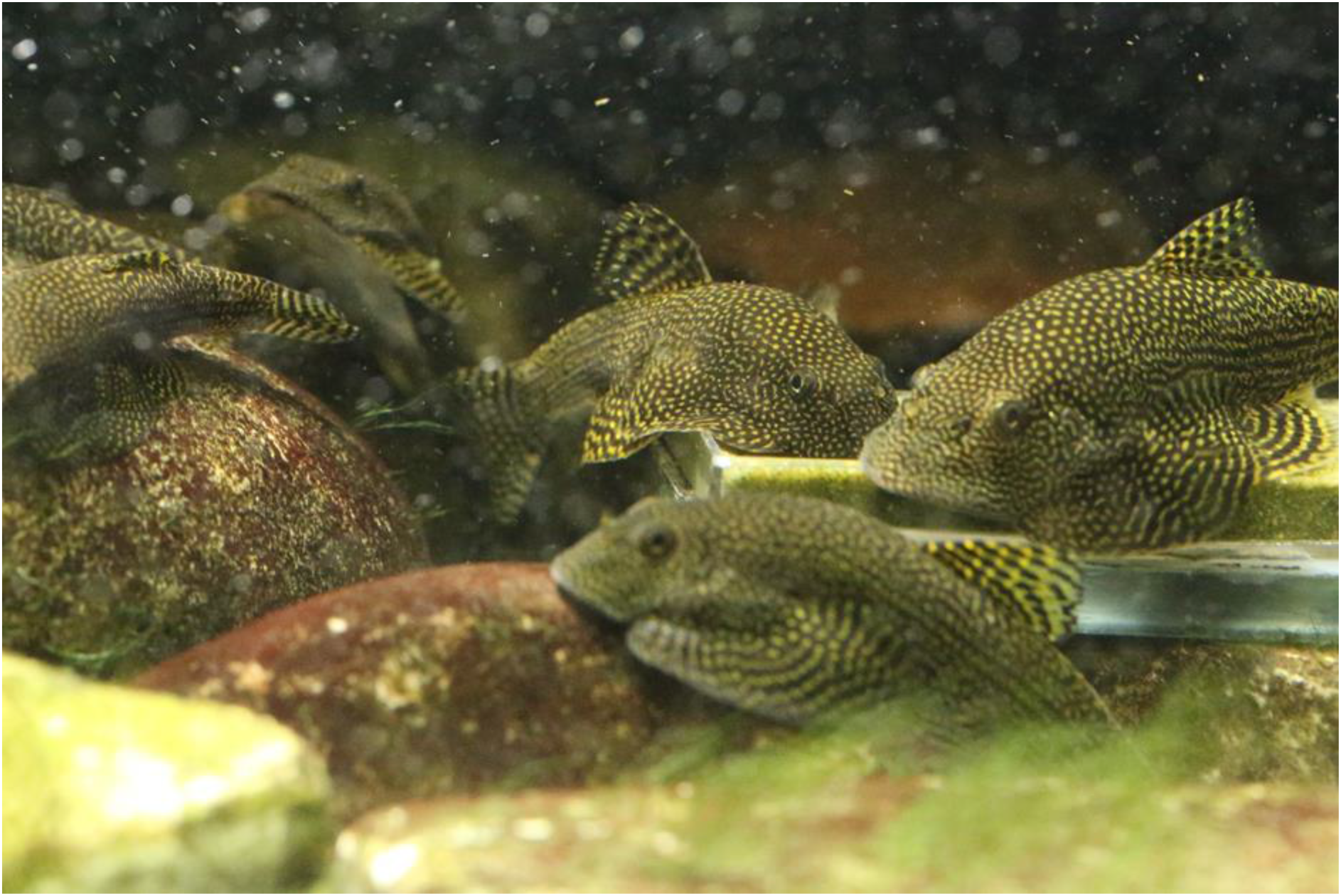
Sewellia ‘SEW01’ Spotted Butterfly Loach used in this study. These fish are aggregating around gel based food. Each are roughly 0.07 m in total length. Members of this species are not territorial during feeding and are regularly in physical contact with each other and almost permanently with the substrates, they are visually attentive to movement in and outside of their tank.

This introduction covers the following subjects in brief summary; 1) distribution and radiation of species, 2) locomotion of loaches, and 3) natural environment and field observations, 4) Functional morphology through experiment, CT (computer tomography) Scans and models of fish.

Hill stream loaches prefer mountainous stream conditions and are not found in the lowland sections of catchments. This pattern of radiation led to the theory that all the hill stream loaches evolved from a limited set of common ancestors that could inhabit both lowland streams as well as the mountainous streams that overlap with the present habitats of hill stream loaches. This theory proposed in the 1950’s (Hora, 1952) is broadly supported by physiologically informed phylogeny (Sawada, 1982). The radiation concept infers that similar physical environments and challenges have shaped these fish; consequently hill stream loaches which are phylogenically distant have remarkably similar morphology, especially those with the most elaborate adaptations to fast flow. For instance members of the genus *Sinogastromyzon* are almost indistinguishable from fishes of the genus *Sewellia* and yet they are in different families (*Balitoridae* as opposed to *Gastromyzontidae*) and endemic to locations that are unconnected mountainous regions and physically distant from one another (Taiwan and Thailand respectively). All these fishes are valuable to humans for the aquarist’s trade and to a lesser extent as a food source. As a result, their distribution has been modified by human activities (see Tan 2006 for example of intentional relocation). In addition, some species are hybridised intentionally or accidently in the fish trade. However they are generally very difficult to breed in captivity and reliable reports of captive breeding in some species (*Gastromyzon*) do not exist (Tan 2006).

Hill stream loaches locomotion is unlike that of open water fishes (De Meyer and Geerinckx 2014, Roberts 1982). Our observations confirm that locomotion fits into three categories, 1) sucker based, 2) ground effect based, and 3) swimming. Here we present video evidence of these modes (Supplementary material). 1) Using only their suckers, without apparently using their caudal fin or tail, or any other water moving fin, they can make movements in any lateral direction with respect to their body orientation; described previously as creeping or crawling (Roberts 1982). This method of locomotion has been defined as a process of sucker release and re-attachment (De Meyer and Geerinckx 2014); we consider this as a hypothesis to be tested in this study. Although De Meyer and Geerinckx (2014) also describe unculi (single cell protrusions - Roberts 1982) on the fin rays which may enhance friction and suggest the alternate hypothesis in which the fins are the main propulsive foci. The combination of requirements for respiration and feeding while remaining attached, or engaged in sucker based locomotion, involves a number of morphological adaptations to control water (pressure) underneath the entire body of the hill stream loach (De Meyer and Geerinckx 2014). In this study we aim to clarify, for the first time, exactly how and with what morphological adaptations, these fish control pressure and water flow under their bodies. 2) Ground effect swimming is a mode of movement whereby hill stream loaches use their tail to provide the main propulsion but glide very close (within 1 or 2 millimetres) to hard substrate, and 3) is conventional swimming in open water which they perform comparatively rarely and for distances of less than 0.5 m, as an escape or startle reaction. (These are illustrated with video sequences – see supplementary material - locomotion).

The crawling mode of movement (type 1 above) has been the subject of recent scientific interest because such movement is thought to be an evolutionary precursor to tetrapod walking on dry land (Flammang et al 2016). In particular the skeletal structure of the walking cavefish (*Cryptotora thamicola*) has been described; this species has developed a pelvic girdle and is able to climb waterfalls in fast flowing water in caves and can walk up wet surfaces in air. The blind cave fish is a loach in the family *Balitoridae* and is endemic to Thailand. It is also listed as vulnerable on the IUCN red list (ICUN 2018). It is protected and is difficult to capture in the wild and therefore a challenging animal to study (Flammang et al 2016). Here we contrast the *Cryptotora thamicola* skeletal structure with the more accessible and captive bred *Sewellia* sp. We also compare the walking gaits. Therefore this work contributes to this field by examining if hill stream loaches may be more convenient or appropriate study subjects with respect to tetrapod gait development than the blind cave fish.

We studied the natural environment of endemic *Gastromyzonid* hill stream loaches at the Danum Valley Field Centre, Borneo. Very little is known about the ontogeny of the Borneo suckers and the eggs and young have never been described (Tan, 2006), nevertheless it is clear from their distribution and our observations that waterfalls over solid substrate represent physical barriers which they climb. However, they are most often observed and captured at a depth less than 1 m, feeding over strongly sunlit algal covered rocks in slower-flowing (< 2 m s^−1^), clear pools (Personal observations (Borneo), also Tan 2006 (Borneo), Yu and Lee 2002 (Taiwan)). Thus climbing represents a fitness advantage – most likely reducing competition by affording access to algae-rich pools which are also less accessible to predators. If this is the case climbing may be an activity which is crucial to life history, and crucial to morphological evolution but which is required or performed relatively rarely. This might lead to behaviour which provides physical conditioning for climbing while in relatively benign environments; a characteristic behaviour of hill stream loaches is sparring between individuals of similar size (Roberts 1982). Here we present video clips of this behaviour in adults and juveniles (supplementary material).

To understand the functional morphology of these species we use techniques that have been developed for measuring the suction power of gobies (*Gobiidae*) (Cullen et al. 2013, Maie et al. 2012) and repurpose them for use with hill stream loaches in order to allow direct comparison. (The *Gobiidae* of Hawaii also climb by the use of suckers (Cullen et al. 2013)). We also examine hill stream loach anatomy using Computed Tomography (CT) scans which allows us to overcome a key problem which happens with sucker fish in general during preservation. The fixation step changes the relative position and orientation of the bone structure. The prominent suckers are activated by ligaments under tension and therefore post-mortem when muscles relax, and the substrate is no longer in a position to hold the suckers extended, the suckers and bodies take up hunched poses that are unnatural or only transient poses taken up by the living fish (for instance see identification pictures in Tan 2006). Therefore to examine the fish skeleton in natural positions it is helpful to separate each bone from its neighbours and manipulate them independently. Micro CT scanning offers internal views of the fish as solid 3D volumes and allows a virtual dissection of the skeleton but may not overcome this fixed positioning problem. However, 3D modelling software such as Autodesk Maya (autodesk.com) and Blender (blender.org) is widely available and this coupled with powerful desktop computers allows for CT scan volumes to be exported as 3D shapes that can be manipulated, moved and printed. The resultant 3D models can also be used in other analysis software such as hydrodynamic modelling or stress modelling (for instance ANSYS (ansys.com)).

## Materials and Methods

### Summary of observations

We took CT scans of several species of hill stream loaches. We made 3D models of sections of the scanned results using 3D animation software in order to dismember the results. This allowed for the construction of solid isolated components that can be manipulated with respect to each other and printed or otherwise processed. We did this in order to observe the physical structure of the skeleton related to the suckers and isolate this from the other structures. We used Frustrated Total Internal Reflection (FTIR) to image the opening and closing of the suckers in living loaches. FTIR photography enhances the parts of a soft damp body which are in contact with a transparent plate surface in proportion to the pressure with which they press the surface (Risse et al 2013). We also made images of fish climbing a vertical glass wall. The pressure under the fish was measured in various positions to create a kind of ‘heat map’ of pressure as it climbs the wall in order to contrast this with the FTIR measurements and produce a quantitative observation of functionality. We measured the drag of a moulded model fish on the same vertical wall. We also took multiple video sequences of behaviour and still frames of anatomy in our experimental tanks which have been designed to mimic the natural habitat of the fish.

### Study subjects

*Sewellia lineolata*, *Pseudogastromyzon* sp., *Homaloptera orthogoniata* and *Gastromyzon punctulatus* were used for the CT scanning and *Sewellia* sp. The ‘SEW01’ Spotted Butterfly Loach (Seriously fish, 2018) is the primary subject for our functional study as we could reliably breed these continually in captivity. This latter species has not been reliably incorporated into the standard scientific nomenclature but is readily available in the aquarium trade and is reliably described as part of the genus *Sewellia*. We also keep *Gastromyzon* sp., *Homalopteroides* sp., *Pseudogastromyzon* sp. in our tanks to inform our general understanding of these species.

### Husbandry

Our aquarium tank design allows water to be pumped to a shallow overhead section and flow back down a ramp and a small waterfall into the main tank. The tank system also provides dark spaces, aeriation and white noise through falling water, and deeper sections of slower moving water. The growth of algae was promoted through strong (500 W) incandescent overhead lighting, nutrient fertilization, high pH (~8.5) buffered by medium to high carbonate hardness (kH > 150 ppm), standard fish food and variable flow conditions. We supplement algae feeding with gel based foods. The substrate is only large smooth rocks which are removed and replaced, in different positions, on a weekly basis. We have kept snails, shrimp and other fish (*Danio rerio*) co-habiting in these tanks in an attempt to replicate the natural habitat.

### Computed Tomography (CT) Scan

Where necessary, fish were euthanized with overdose of MS-222 (500 mg/L). They were then fixed overnight in a solution of paraformaldehyde 4% in Phosphate Buffer Saline at pH 7.5 and moved to a 70% ethanol solution for long term storage. We used the GE v|tome|x M 240 kV microCT system (www.gemeasurement.com) at the Hounsfield Facility, University of Nottingham. *Sewellia lineolata* was scanned at a resolution of 35 microns, with X-ray tube settings of 100 kV and 200 μA collecting 2785 projection images over a 360° rotation of the sample. *Gastromyzon punctulatus* was scanned at 18 micron resolution, with X-ray tube settings of 85 kV and 180 μA collecting 3203 projection images over a 360° rotation of the sample. Projection images were reconstructed used Datos|x REC software (www.gemeasurement.com). The images were initially analysed and manipulated in VGStudio Max and viewed in MyVGL software from Volume Graphics (www.volumegraphics.com). The software was used to output the files in STL format.

### 3D software manipulation and printing

We filled in holes in disassembled polygon volumes with manually added triangular elements. Models were prepared for printing using Autodesk NetFabb (www.autodesk.com). The models were printed using a variety of commercial printing companies and techniques – for the skeletal structures we found Stereolithography (SLA) 3D printing using Formlabs Form 2 (www.formlabs.com) the best option where the models were strong enough to handle and not brittle, although the finished size was limited and the models required considerable finishing to remove the remains of the support structures.

### Wall climbing

Fish were encouraged to climb through a passive voluntary response rather than a positive or negative artificial stimulus (Fig 2).

**Figure 2.**
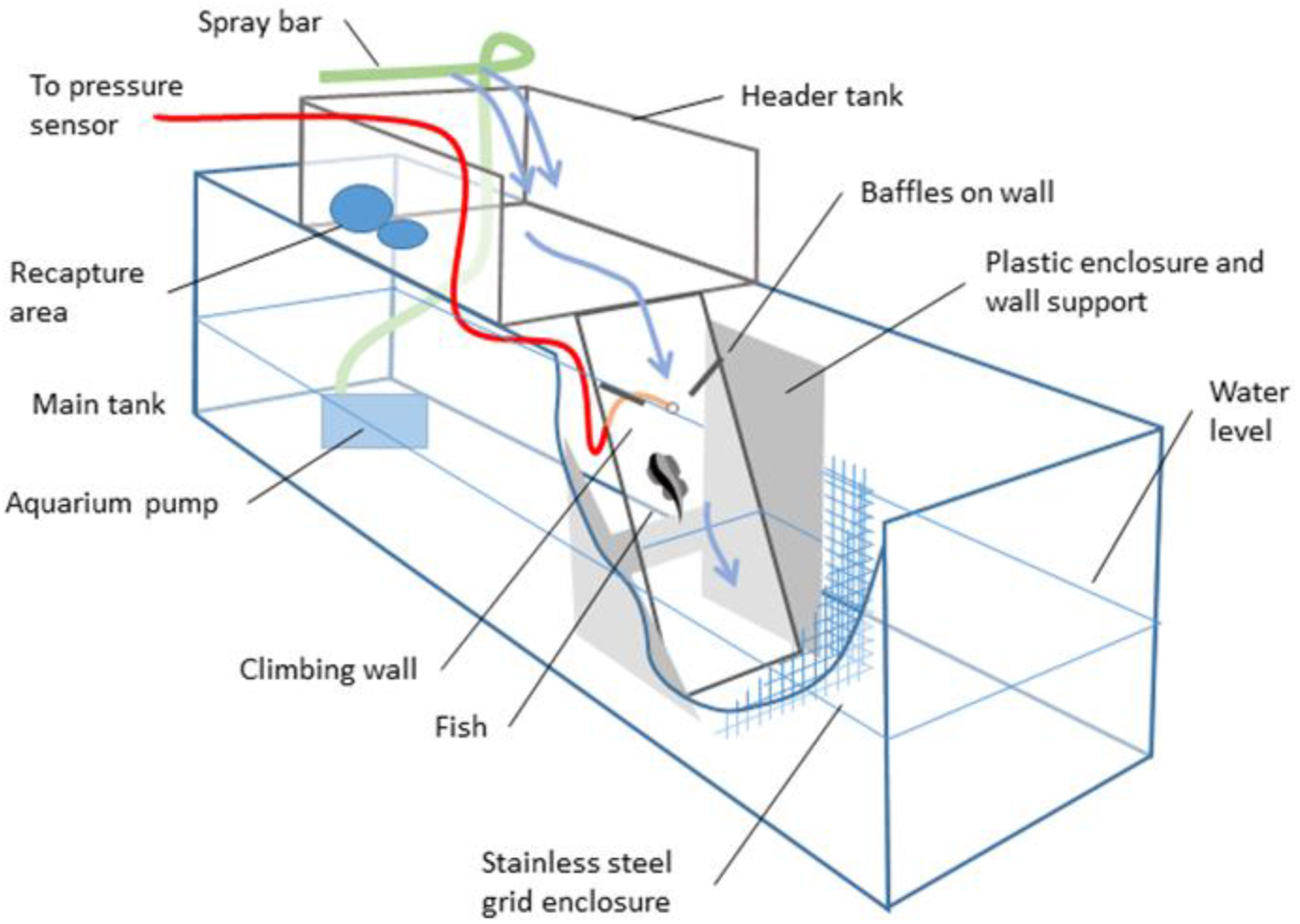
Wall climbing aquarium. The tank is a standard 0.006 m float glass aquarium tank, 0.9 m × 0.4 m × 0.3 m. Depth in the main tank - 0.15 m, and waterfall surface 0.01 m float glass at an angle of ~10o to vertical. Water was pumped from the lower tank to a header tank distributed through a standard aquarium spray bar using a submersible aquarium pump rated at 1000 litres per hour. All the pumped water returned to the main tank via the waterfall. The refuge area was a plastic tub in the header tank with an opaque lid with smooth rocks as shown.

We placed baffles on the face of the waterfall to shape the water movement over the face in order to encourage climbing on a route that passed over the sensor. The fish would usually voluntarily climb the wall, but sometimes did not and we moved them by hand to the refuge area after about 20 minutes if that was the case. We placed multiple fish in the apparatus together. The refuge area at the top of the waterfall was a plastic tub which was covered with opaque black material. Any fish showing signs of stress (e.g. detachment and attempted open water swimming, or attempted attachment to the stainless steel gridded surfaces and/or rapid and sustained fluttering of the posterior pectoral fin margin) would have been immediately removed, however that did not occur. All the fish that were involved in the climbing observations were checked regularly afterward and no change in behaviour or condition was observed for the following 30 days.

### Pressure sensor

A single hole of diameter 2 mm was drilled through the 10 mm glass plate used for climbing (Fig. 2), and expanded to a 4 mm hole on the underside. A silicone tube of internal diameter 2 mm and external diameter 4 mm was inserted into the rebate and aquarium silicone sealant used to secure it. The tube was 0.3 m long and connected via a stainless steel barrel connector to a similar silicone tube 0.05 m long which had been glued over the top of a pressure sensor (Adafruit BMP280 Barometric Pressure + Temperature Sensor Breakout – www.adafruit.com). The silicone tube glued to the breakout board totally enclosed the BMP280 chip and was sealed with epoxy resin (www.go-araldite.com). The breakout board was connected to an Arduino UNO (www.arduino.cc) and this was connected to a PC running MATLAB (www.mathworks.com). The Arduino was operated using a standard script for the BMP280 breakout and MATLAB set up to parse information returned by the Arduino on the serial interface. A GUI (graphical User Interface) was written in MATLAB to display the output of the pressure sensor in real time as a graph and save the output on the PC. The Arduino also controlled an LED (Light emitting diode) which was used for timing synchronisation as it was placed in a position visible to the video camera which was recording the fish movement. The LED could be operated via a button on the pressure recording GUI and its operation caused a false signal and a suspension of the pressure results for 1 s. The false signal and suspension on the pressure output results could be matched to the light on the video to synchronise the time in another GUI used for analysis.

All analysis of pressure sensor results was performed in MATLAB using a custom GUI which ran the video and pressure results graph together. A template of a fish outline was translated, rotated and scaled on each video frame to best match the outline of a fish interacting with the area in which pressure recordings were made. The manual classification of fish position was more convenient than automatic classification as the outline of the fish was broken up by the chaotic water surface (see supplementary material for video example).

### Load Cell

A load cell was used to measure the drag of a model fish on the waterfall. The load cell was firmly attached to a ‘Z’ shaped cantilever arrangement of two beams of carbon fibre reinforced sheeting (0.15 m × 0.01 m × 0.006 m) and calibrated with a known weight. The load cell was a 100g Micro Load Cell (www.robotshop.com) which was connected to an Arduino UNO (www.arduino.cc) via an HX-711 amplifier (AVIA Semiconductor www.aviaic.com) and a standard load cell script used. The Arduino was connected to a PC running MATLAB and a GUI used to graph the parsed output of the serial line. The load cell beam was attached to the moulded model fish via nylon fishing line (0.24 mm 4.5 kg, www.drennantackle.com). The load cell data could be synchronised with video data of the situation in the same way and using the same GUI as the pressure data above.

### Frustrated Total Internal Reflection (FTIR)

We used an 8 mm thick sheet of transparent acrylic plastic. Along one edge we drilled 12 × 6 mm holes in position to accommodate the 12 protruding LEDs embedded in a submersible decorative bubble curtain device (manufacturer unknown). We attached reflective plastic film to the opposing edge and as a 0.025 m frame around the edges on both faces of the sheet to act as waveguides near the emergent and reflective boundaries. All other areas where light was directly emitted were covered by multiple layers of duct tape. The FTIR device was used in two methods, 1) as a nearly horizontal plate in total darkness, with a high speed camera (Sony RX10 II see following section) positioned directly below running at 250 frames per second (fps). The plate very gently sloped toward the edge of a normal husbandry tank and overhung the edge by several centimetres. The fish were released in the centre of the plate in very shallow water and encouraged to move to the edge of the plate by being softly touched by hand, and they voluntarily dropped back into the tank. Exit from any other side of the plate was blocked, but the fish always travelled in the direction of the very slowly flowing water down the slope to the tank. This is normal behaviour for these fish which often voluntarily move into very shallow water (~1mm depth) from, and back to, a main deeper pool. 2) The same FTIR device (cut down in size to 0.3 m × 0.1 m) was used as a replacement for the waterfall face in the climbing experiments described above and the fish movement was filmed from underneath the plate – there was unavoidable ambient light in this case due to the very active water surface.

### High speed photography and analysis of video

In general we used a Sony RX10 II bridge camera (www.sony.com) which has a nominal maximum video capture rate of 1000 frames per second, we shot video from 30 fps to 1000 fps depending on the required resolution and the available lighting. We also used a GoPro Hero 4 (www.gopro.com) for filming the FTIR device in the climbing wall when the camera was in a semi-submersed position behind and under the wall. For still photography we used a Canon EOS6D with a Canon EF 100 mm Macro lens and a Canon MR-14EX II Macro Light Ring (www.canon.com). For tracking the movement of images in video frames we used point trackers in the computer vision system toolbox of Matlab (www.mathworks.com) using default parameters.

### Moulded model fish for calibration and drag measurements

We used a two part silicone moulding material (www.smooth-on.com) to cast a soft mould of a dead *Sewellia lineolata* (0.053 m Total Length (TL)), we used this mould to make multiple life-size replicas from a harder version of the same silicone material (Smooth-Cast^®^ 300) of the fish with a flat ventral surface. The reproduction of surface detail was good to a resolution of about 0.1 mm (confirmed by optical microscope), the weight was similar to an actual wet fish and the body negatively buoyant and sank at approximately the same rate as a natural wet dead fish. These were used in the calibration of the climbing wall experiment in two ways: 1) explained above using the load cell to measure total drag plus bodyweight of fish, and 2) as a calibration of the pressure response to a non-sucking fish shape with a flat ventral surface. There was random normal variation in the pressure instrument readings and some drift over longer time periods and so the moulded model was used to determine a low bound on what pressure could be reliably recorded during an interaction.

## Results

### CT Scan

The scan results show the elaborate morphological adaptions of *Sewellia* and *Gastromyzon* species when compared to typical open water fish (Flammang et al 2016). In particular the puboischiadic plate is the most prominent skeletal feature on both species (Fig 3). The pelvic girdle is similar to the cave fish (*Cryptotora thamicola*) (Flammang et al 2016) in that one rib (called the pleural or sacral rib) is hypertrophied and forms a closed loop with the puboischiadic plate. This pleural rib is described by Sawada (1982) as being connected to the puboischiadic plate by a strong ligament (invisible in CT scan) in closely related species.

**Figure 3:**
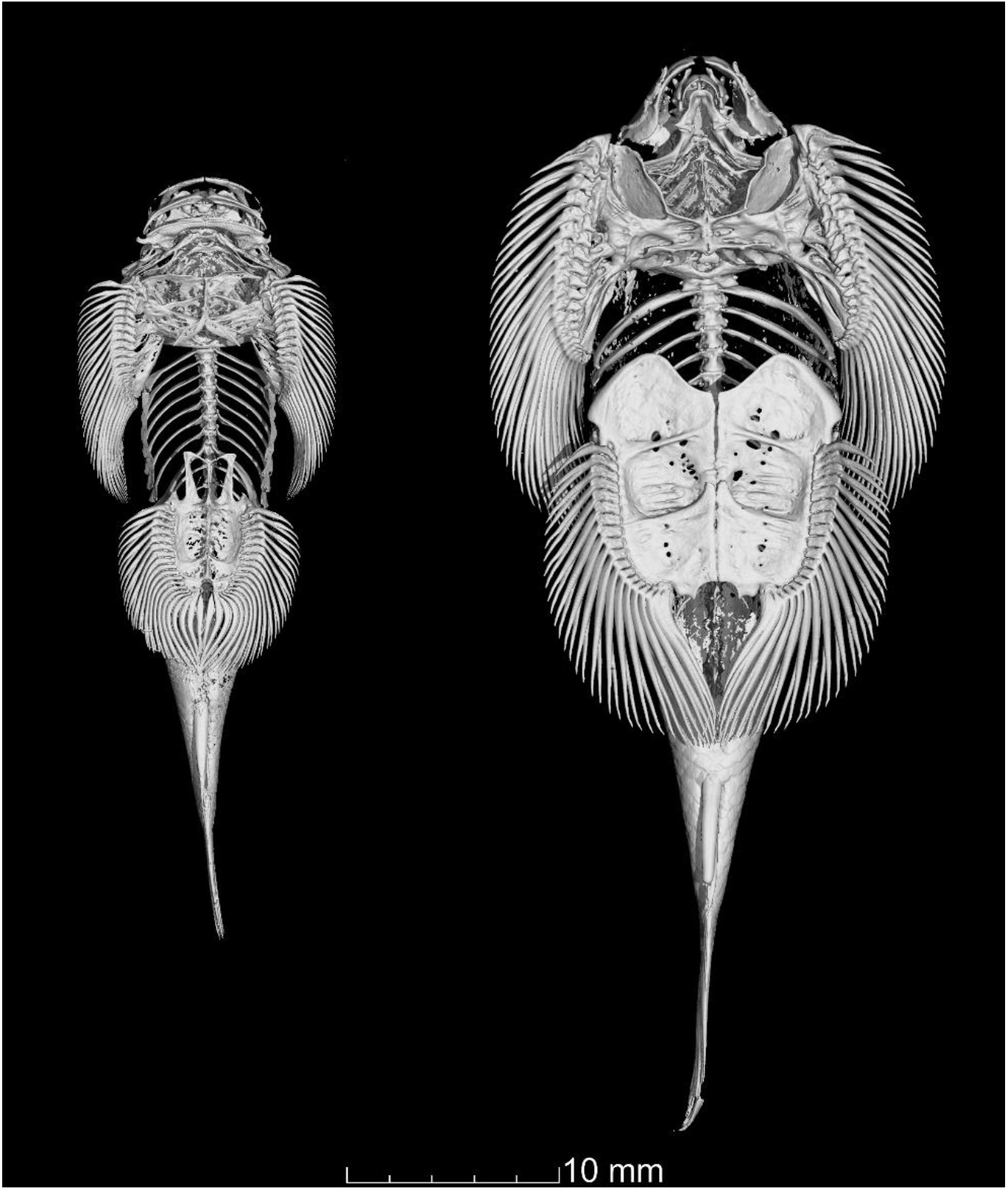
CT scans of Gastromyzon punctilus (left) and Sewellia lineolata (right). The Sewellia appears more elaborately adapted for suction climbing with a more prominent pelvic plate and distinctive ‘shoulder blades’ of the pectoral girdle. Both fish have well developed sucker margins of elaborately adapted fin rays. One major difference between these two species is the attachment position of the pelvic puboischiadic plate. On the left the pelvic girdle including pleural rib meets the plate anterior to the fin ray attachments, whereas on the right it is at the posterior of the plate – this dramatically changes the position, major caudal muscular attachment and size constraints of the plate.

The fin rays in the *Sewellia* are more numerous and more complex in shape than either the *Gastromyzon* or the *Cryptotora* (but less in number and degree of adaptation than the *Sinogastromyzon* (see Sawada 1982)). In both *Sewellia* and *Gastromyzon* the pelvic plate is flat and forms a single contiguous surface. The entire pelvic plate is freely articulated with respect to the body with attachment axis defined by the strong stiff but flexible joint with the sacral rib (Fig 4 Panel A). Thus the pelvic sucker is aligned at a tangent to the spine. Whereas the pectoral sucker is made of the head shield (modified cliethrum) and two sets of three articulating bones (modified radials) on each lateral posterior section of the pectoral sucker areas (Fig 4 Panel A). These articulated redials are used in pectoral fin fluttering which is a characteristic behaviour of hill stream loaches (De Meyer and Geerinckx 2014) and suggest that this is not effected via movement of the fin rays alone. Overall the pectoral sucker area is more skeletally complex and more embedded into the main body structure than the pelvic sucker. Other interesting features are the relatively small gill area when compared with open water fish and the Weberian capsules, which are elaborately adapted in all the hill stream loaches including the *Homaloptera* (see supplementary material). Weberian capsules are thought to be hearing organs adapted from the swim bladder in many benthic fish which require negative buoyancy. All hill stream loaches examined by us are negatively buoyant and do not have swim bladders.

**Figure 4:**
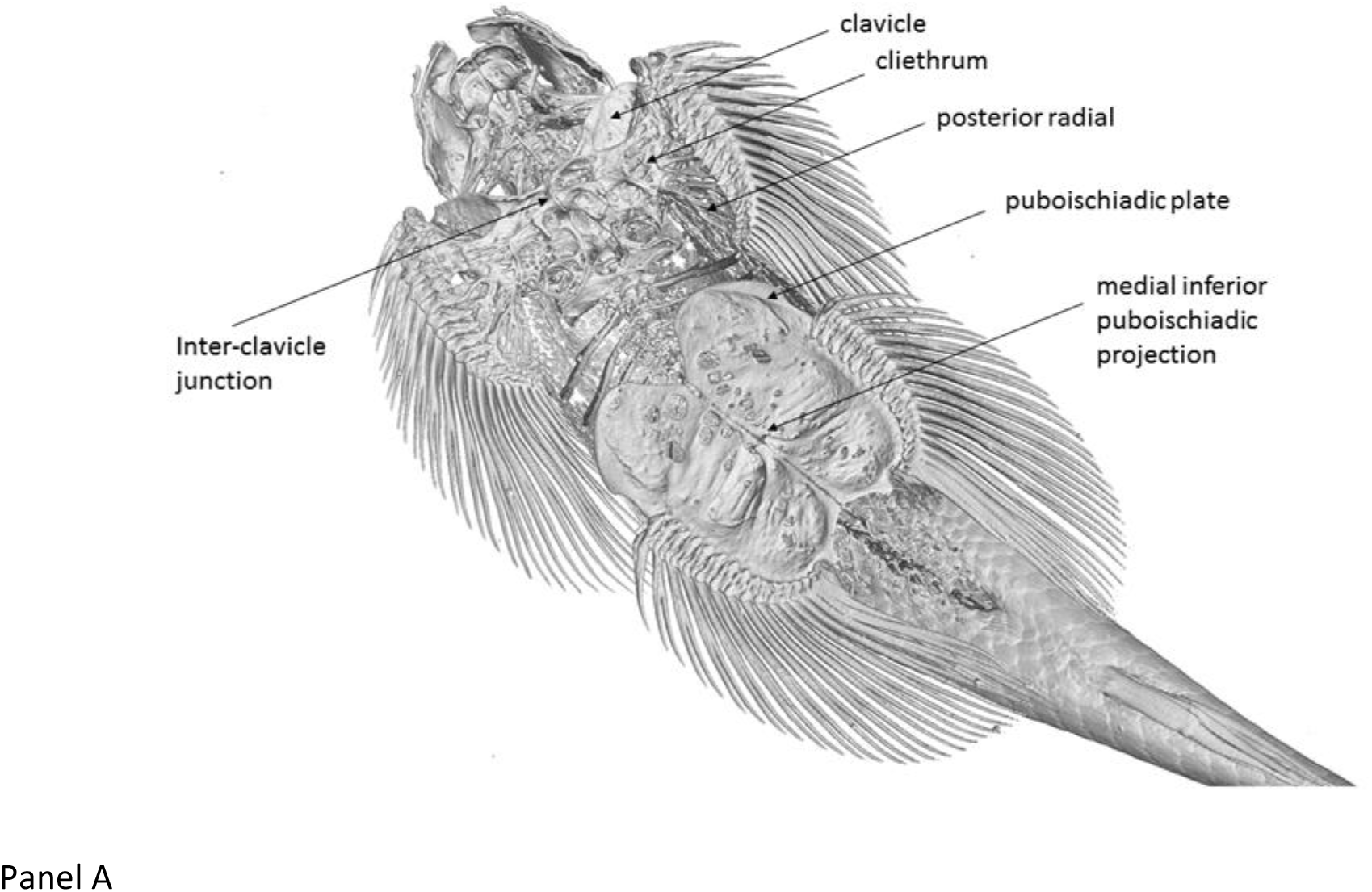
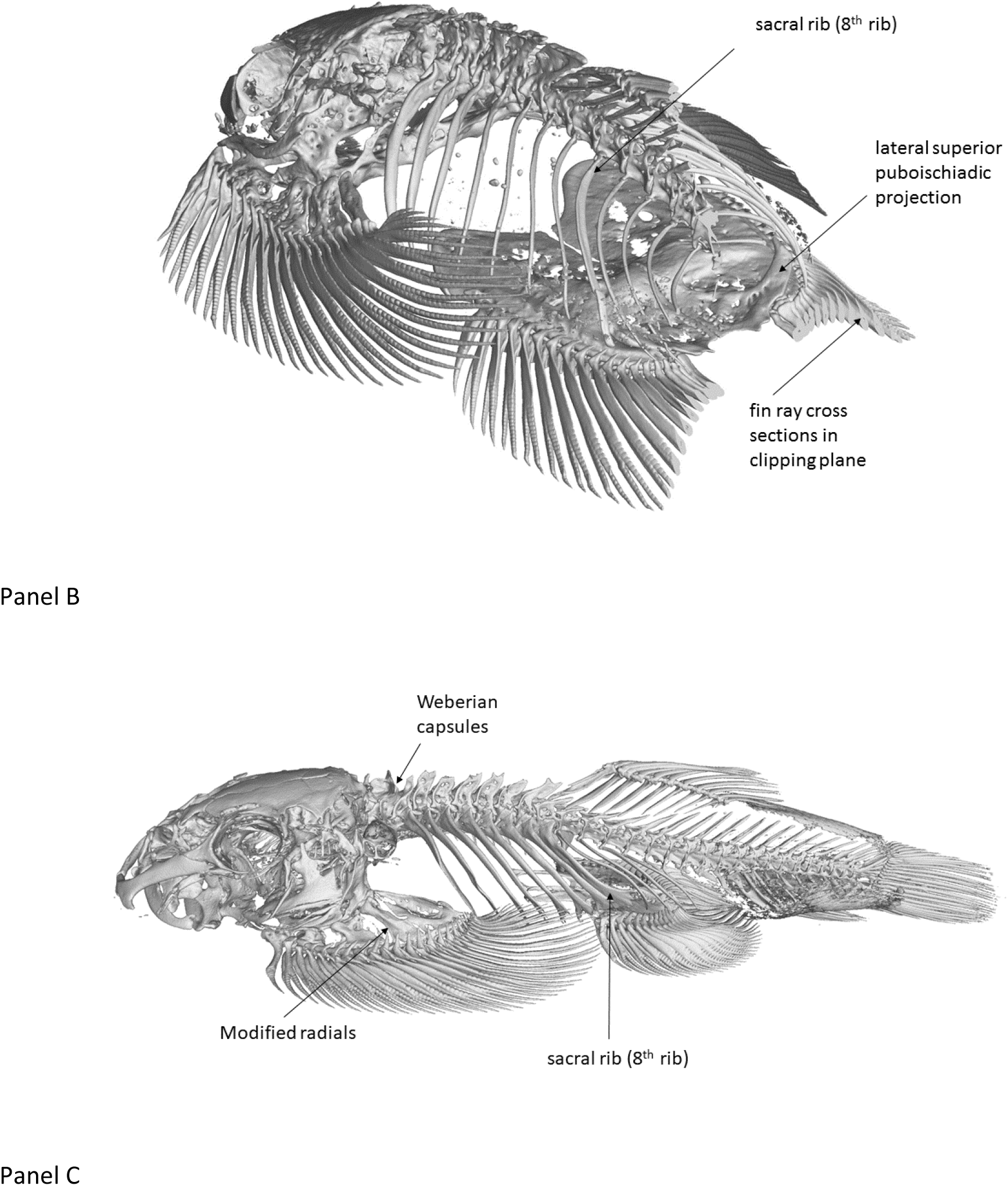
CT scans of Sewellia lineolate and Gastromyzon punctilus. The projections are 3D perspectives so the scale varies but a guide is that the puboischiadic plate is about 0.01 m wide at its widest perpendicular to the spine. Panel A: Ventral view. The fish is in a natural pose of lateral curvature. The head, clavicle and cliethrum are aligned with the anterior of the spine which forms a smooth curve throughout its length, whereas the pelvic plate structure is aligned with a tangent to the spine and pivots at a point behind the medial inferior puboischiadic projection at the position of attachment of the pleural/sacral rib. The medial projection and medial ridge just posterior to it are positions at which the ligaments that form the pelvic sucker are attached, the other ends of the ligaments are attached to each of the processes on the medial ends of the fin rays. In the case of the pectoral sucker the ligaments from each of the fin rays attach to the inter-clavicle junction. Panel B: Dorsal view. The 3D volume has been sectioned by a clipping plane to remove the tail from view. The first three ribs and the sacral rib are strengthened in comparison to the others. The sacral (pleural) rib is not fused with the puboischiadic plate but ends very close to it at the base of the lateral superior puboischiadic projection forming a strengthened arch with the spinal vertebrae at the apex. The lateral superior puboischiadic projections are highly buttressed horn shaped raised processes and are likely to be the primary anchor for the pelvic sucker with respect to the main caudal musculature of the fish. In this specimen the sacral rib in the foreground has been fractured in life about ¾ along its length, the bones have fused slightly. Panel C shows the lateral view of Gastromyzon punctilus, the broad calcified processes at the distal end of the plural ribs are typical of many of the balitorid and gastromyzonid fish including the blind cave fish and are thought to be the main points of muscular attachment to the pelvic girdle, which is dissimilar to the Sewellia above. The Weberian capsules are thought to be hearing organs and are common among bottom dwelling fish including all the fish in this study.

The one major difference between the *Sewellia* sp. and the *Gastromyzon* sp. is the attachment of the sacral rib to the puboischiadic plate. In the *Sewellia* sp. the sacral rib attaches to the posterior of the plate posterior to the attachment of the fin rays whereas in the *Gastromyzon*, *Cryptotora* (Flammang et al. 2016), *Pseudogastromyzon* and *Homaloptera Orthogonata* (Supplementary material) the sacral rib attaches at the anterior of the plate and to the anterior of the fin rays. In all the latter cases where the sacral rib attaches to the anterior of the plate the sacral rib is hypertrophied, heavily calcified and forms a broad flared process at its distal end (most elaborately in the *Cryptotora* (Flammang et al. 2016)) (see *Gastromyzon* in Fig 3 Panel C). Flammang et al. (2016) assumed this was an indication that the main caudal muscle attachment was on the rib. In contrast the *Sewellia* sp. does not have this flared process and the sacral 8^th^ rib is hypertrophied but not proportionally as much, or as variably along its length, as in the other species (Fig 3). However the *Sewellia* does have two major skeletal protrusions on the puboischiatic plate which are absent in the other species; the lateral superior puboischiadic projections (Fig. 3 Panel B). So we assume that the major caudal muscular attachment in the *Sewellia* sp. is to the plate rather than the rib. Then the plate is anterior to the musculature attachment rather than posterior to it, and this may have major ramifications in terms of the subsequent development of the two groups which differ in this respect (Fig. 3).

### 3D models derived from scans

The 3D models help understand the way in which the pectoral girdle both encases the back the head in a bowl-like shape (mainly the cliethrum) but also provides buttressing and structure to the anterior base of the pectoral sucker (Fig 5). It appears that the head may be free to move independently but Sawada (1982) notes that in homalopterine fishes (which includes all the fish in this study) the skull, pectoral girdle and Weberian capsule (which is also particularly well developed in all the fish in this study) are joined by extensive connective tissue into a rigidly combined unit (Sawada 1982 pp. 137). Physical examination of dead specimens and observations in captivity show that for *Gastromyzon* and *Sewellia* the head has free movement, but of limited extent, in the ventro-dorsal plane (nodding up and down) but little or none in the lateral plane. This may be a particular adaptation to feeding and breathing while attached to a surface related to the positive pressure that can be maintained under the head which is described in the following results (see De Meyer and Geerinckx (2014) for an in depth discussion of feeding and breathing). Therefore the structure of the cliethrum may be related to rigid support of the pectoral sucker system and isolation of the skull and breathing chamber (buccal cavity) from suction pressure effects caused by the pectoral sucker. Parts that have been disassembled and rebuilt into individual polygon volumes were printed and examined in groups and individually.

**Figure 5:**
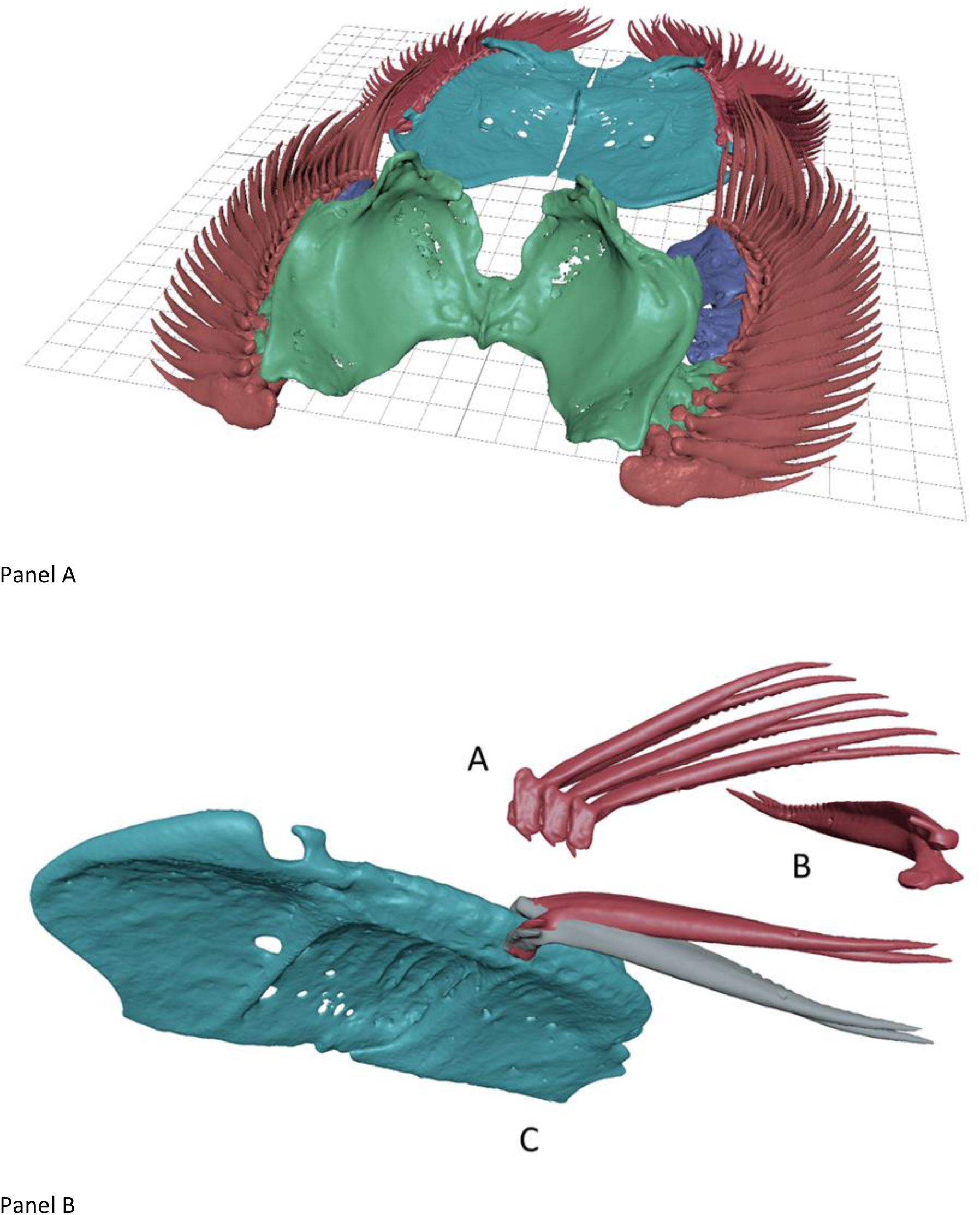
3D Modelled skeletal sections. Panel A: Pelvic and pectoral sucker bone structures of Sewellia lineolata from micro CT scan transferred to 3D modelling software as solid volumes. The individual fin rays are coloured red, the pelvic plate teal, the fused clavicle, cliethrum and other bones of the pectoral girdle are green, and the mobile radials of the pectoral girdle are purple. These volumes can be 3D printed or used in other software such as hydrodynamic modelling software and for strain analysis. We used them in this study to examine the likely movements of one set of bones in relation to another. Panel B: Group A is a ventral view of 3 fin rays. The individual fin rays interlock with their neighbours due to a tab like projection on the ventral surface. This prevents any one ray moving dorso-ventrally with respect to its neighbour and locks the rays as a sheet, it allows a fan-like movement in the anterio-posterior orientation. Group B shows a ray in an orientation to highlight the extended sharp crest of the ray in the dorsal direction, this stiffens the ray in this orientation and leads to asymmetric flexibility of the ray. Section C shows one lateral half of the puboischiadic plate in teal in ventral view, the fin rays in red. The grey fin ray is a duplicate of the red. There is a ridge in the puboischiadic plate which acts a brake on the potential movement of the fin ray and this is matched on the dorsal surface. These files in 3D printable. STL format are published in supplementary material.

### Wall climbing

The fish held position on the vertical glass wall under the waterfall for long periods of up to 5 to 8 minutes between intensive bouts of climbing that typically last no more than 2 or three seconds (See supplementary material videos). These times are consistent with classification of fish activities such as burst swimming as opposed to coast-burst and sustained swimming (maximum sustained swimming rate) (Videler and Weihs 1982). On the occasions where the water was not flowing strongly over the surface of the climbing wall in our apparatus the fish did not climb or rest on the surface, but immediately resumed climbing when flow was restored.

We recorded 1140 pressure data from 17 fish in 17 interactions with the recording point on the wall. An interaction was defined as a sequence when a fish moved over the hole voluntarily – in all occasions the fish was vertically aligned with its head at the highest point. The fish varied in total length between 0.0527 m and 0.0725 m (mean: 0.0634 m interquartile range: 0.0014 m).

The pressure distribution under the fish showed a distinctive pattern of positive pressure under the head and negative pressure (suction) on the rest of the body and fins (Fig. 6). In general the highest suction points were around the pelvic fin margin, and in the gap between the pelvic and pectoral fins close to the sucker margins. There was no significant correlation between the pressure measurements and total length of fish for the head (N = 201, R^2^ = 0.01, p = 0.16) and pelvic (N = 285, R^2^ = 0.11, p = 0.054) regions, but there was a weak but significant negative correlation between the total length and pressure generated in the pectoral region (N = 204, R^2^ = −0.31, p<0.0001). That is; larger fish had mildly higher suction forces on average in the pectoral region but otherwise there was no difference in pressure related to size of fish.

**Figure 6:**
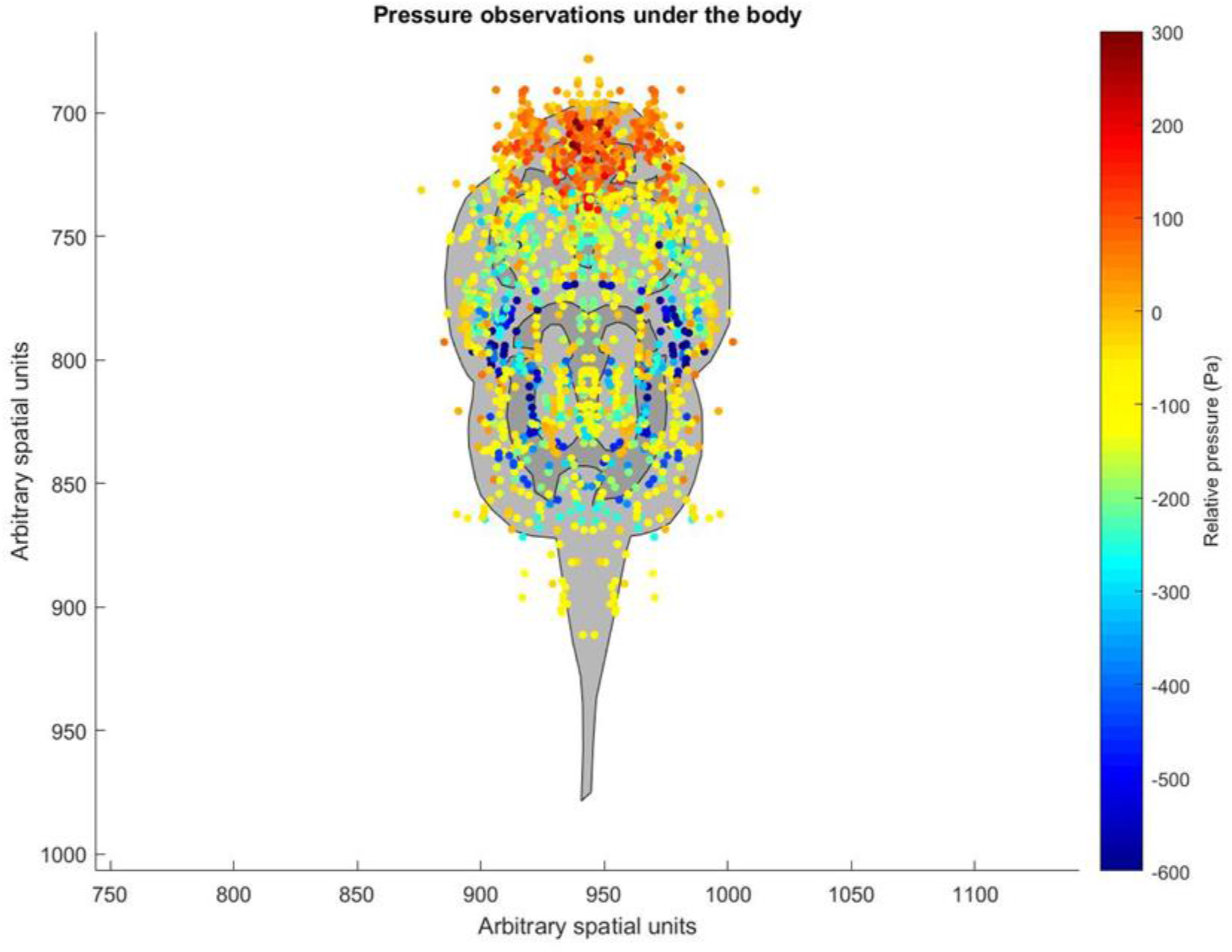
Pressure observations on the ventral surfaces of multiple Sewellia ‘SEW01’ as they climbed a steep glass wall under a waterfall. The colours relate to relative pressure over the baseline pressure without the fish present in Pascals (Pa), 100,000 Pa is roughly normal atmospheric pressure. The area of the head has predominantly positive pressure whereas the body overall had a negative (suction) pressure. The data have been reflected on both sides of the centre line to provide a consistent symmetrical view.

Practically all the pressure recordings under the suckers are negative; indicating that the areas were under constant mild suction throughout the movement cycles. The alternative is true for the head which was usually under positive pressure (Recordings in head area: N=201, mean = 53.6 Pa, standard deviation = 91 Pa, Number of recordings under zero = 43, distribution: mono-modal). The highest positive recording of pressure were near the mouth of the fish.

### Drag measurements and other calibration on the pressure wall

#### 1) Pressure response of a model fish

We used a moulded model fish freely hanging in the water flow which was pulled over the pressure sensor. The moulded fish mass was total length 0.065 m, weight 3.7 g. When the moulded model fish was moved over the pressure sensor we consistently recorded a small positive pressure (~18 Pa) throughout the whole body plan of the model fish. This measurement was confirmed as significant by using a t-test between baseline pressure readings and interaction pressure readings in four interaction events (t-stat = 18, df = 168, confidence interval 16-20, p<0.0001).

#### 2) Pressure of a silicone sucker

A 0.025 m diameter silicone aquarium sucker caused a reasonably constant suction pressure when dragged over the pressure sensor hole; a negative pressure around 5000-7000 Pa on each interaction. A rebound positive pressure spike of up to 200 Pa was evident after the sucker was moved off the sensor hole, which existed over base pressure for less than 0.2 s (one reading interval).

#### 3) Drag of a model fish

The drag measurements of a moulded model of a dead fish on the glass wall were made using the load cell calibrated in grams force (g). N = 20 measurements were made at 4 positions: 1) just under water at the base of the waterfall (mean 2.2 g +/− standard deviation 0.6 g), 2) at the base of the waterfall with entire body out of water (11.6 g +/− 1.7 g), 3) pectoral girdle on pressure sensor position (9.4 g +/− 1.3 g) and 4) at top of waterfall with nose in line with top of climbing wall (5.1 g +/− 3 g).

### Frustrated Total Internal Reflection (FTIR)

#### 1) Horizontal surface

Horizontally inclined recordings of FTIR were made on a video sequence of a *Sewellia* sp. moving in response to a physical stimulus (a touch from a finger). The fish moved away from the stimulus and made around 5 cycles of movement (strides) before coming to rest. The movement was measured by using point trackers. Point trackers automatically identify features in a video frame and attempt to track these points throughout the sequence of subsequent frames. We focus here on one set of movements analogous to two steps in a walking gait that took 34 frames of a 250 fps video. The results identify that the pivots of the movement are on the fin rays rather than the suckers, and so the suckers slide during all parts of the motion (Fig. 7). Qualitatively it can be seen from the coloured figures of the cumulative movement of the points that the gait is a diagonal-couplets lateral sequence gait, which would imply a standing wave on the spine. This is confirmed quantitatively using the point tracker results (Fig. 8 Panel A).

**Figure 7:**
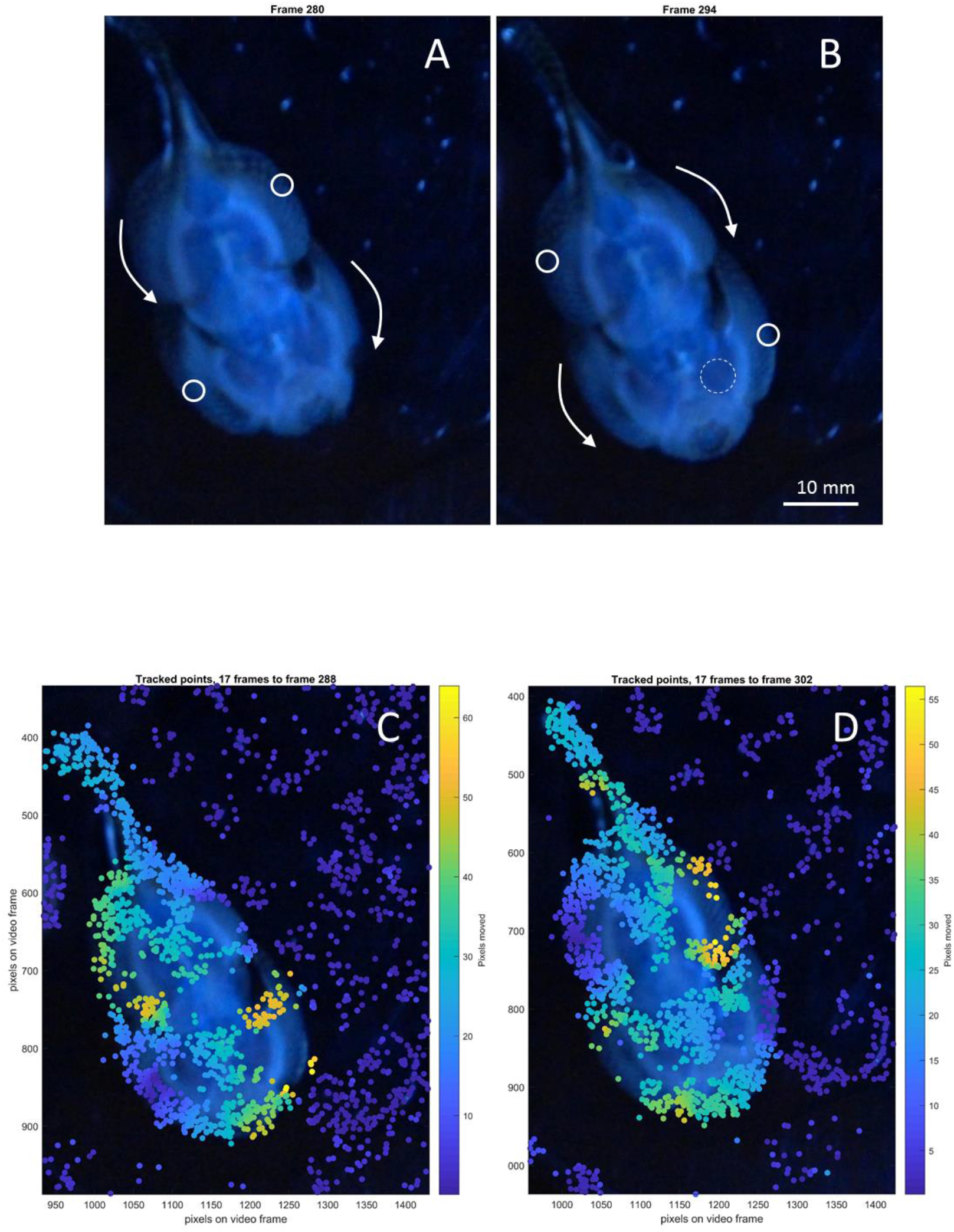
Frustrated Total Internal Refection – Horizontal surface. Panels A and B: Two frames (280 (A) and 294 (B)) from a Frustrated Total Internal Refection (FTIR) video (frame rate 250 fps) of a moving Sewellia sp. Fish. The fish is on a horizontal transparent acrylic plate (8 mm think) and the images are taken from below the plate. The water depth is ~2 mm. The areas where the fish was in contact with the plate are illuminated in approximate proportion to the strength of the contact. The images show the fish midway through two cycles of movement, analogous to two sequential steps in a walk. The fish moves by a twisting action pivoted around two fixed centres indicated by white solid circles. This shows that the fish, in this case, was pushing forward off the fin rays rather than anchoring by a sucker. The suckers continually slide over the surface. The dashed circle indicates the area of a sucker used to determine the change in irradiance during the movement which can be used as a proxy measurement of pressure (see following figure).Panels C and D: Frames (288 (A) and 302 (B)). Tracked points have been indicated by coloured dots, and the cumulative movement in pixels of each dot over the preceding 17 frames (0.068 s) is indicated by the colour. The scale is 10 pixels to the mm. The tracked points show that all four suckers moved throughout the step whereas there were two fixed pivot areas on the fins (top right, bottom left on left panel, and top left, bottom right on right panel – where the bottom fins are pectoral in these views and top are pelvic). The frames in the panels above are mid-way through these sequences and indicate the areas of movement and stasis derived from these point trackers as circles and arrows.

**Figure 8.**
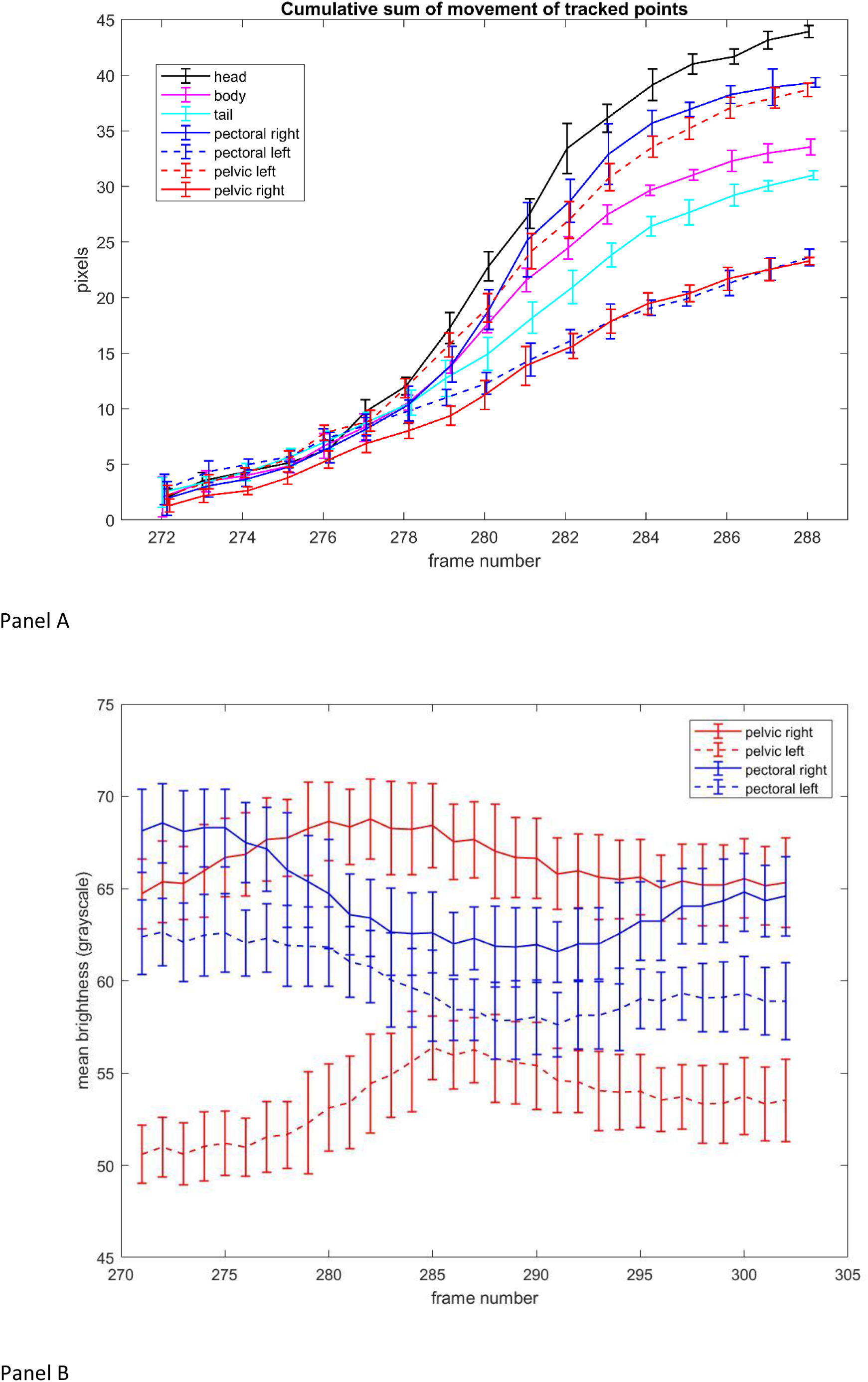
Movement of tracked points during horizontal Frustrated Total Internal Reflection filming. The cumulative totals of movements of groups of tracked points through 17 frames of a video sequence of Sewellia sp. moving over a horizontal acrylic plate. The tracked points were grouped into their initial body positions. The video was running at 250 frames per s, frame spacing in time is 0.004 s, the distance scale is 10 pixels to the mm. The error bars represent one standard deviation of the movement of each group during each frame. The total number of points initialised in each area ranged from N=107 to N=237; points which did not track sufficiently well from frame to frame were dropped (those that were not dropped are shown in the above fig. 11). The figure shows that the pectoral right and pelvic left are statistically inseparable at the end of the cumulative movement and moved significantly further overall than the pectoral left and the pelvic right, which are also inseparable. Both sets are inseparable throughout the sequence and so this demonstrates that in this stride they are in phase. These results are indications of a balanced diagonal-couplets lateral sequence gait. The head moved the most and the tail and body moved at an intermediate rate between the two fin pairs which suggests that the maximum amplitude of the standing wave on the fish’s body was just posterior of the pectoral girdle area. This is extremely unusual for a swimming fish, where the wave on the body is usually a travelling wave and the minimum amplitude is at the head and pectoral area. Panel B: Mean irradiance of pixels within circular areas manually positioned within each sucker area on each frame during the movements of a Sewellia species fish on a horizontal surface while filmed using FTIR. The frame numbers are consistent with the above figures showing individual frames. The designation of left and right is relative to the observer’s view of the above pictured frames. The traces are smoothed with a boxcar filter (extent 7) and the error bars are one standard deviation of values within each filter segment. Higher proximity or pressure causes higher irradiance. The pelvic and pectoral traces are approximately sinusoidal and out of phase, pelvic to pectoral, and in phase right to left.

FTIR as a proxy for applied pressure. The irradiance of an FTIR image is proportional to the proximity or pressure of the overlaying object on or to the upper surface of the substrate; an area which is pushed down hard has a higher irradiance than the same area under less pressure. Higher positive pressure is lighter, higher suction is darker. We measured the irradiance of a circular region manually placed inside each of the four suckers in each frame (Fig. 8 Panel B). The results show a consistent pattern of pressure change for each of the four suckers during the movement cycles.

#### 2) Vertical surface

The glass vertical climbing wall was replaced with an Acrylic sheet similar to that used for FTIR on a horizontal surface and filmed from the other side to the one climbed by the fish (Fig 2). The FTIR images in this case were less distinct in terms of identifying areas that were in contact at various pressures with the surface of the Acrylic sheet because there was higher ambient light and the water surface was turbulent which caused further light interference (Fig 9) but the function of the flutter channels was very well highlighted and the contact pressure pattern of the other anatomical features was confirmed (head, suprapelvic flap, and sucker margins). The video results show that the pelvic sucker group has a wider seal and swings out at a wider angle during climbing.

**Figure 9:**
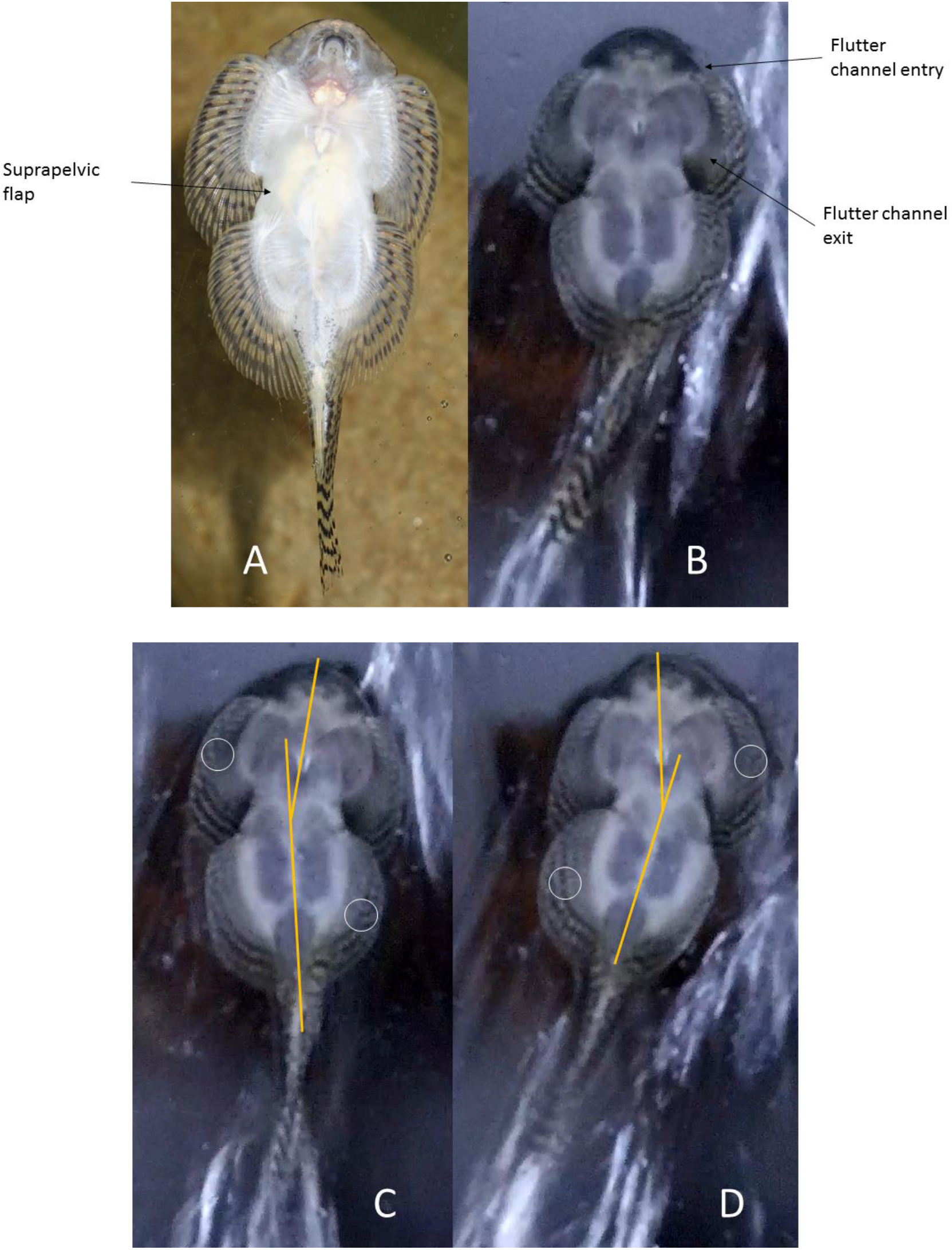
Ventral views of vertical climbing. Panels A and B: Contrasting ventral views of two Sewellia species fish attached to transparent surfaces. The left panel shows a fish photographed using a flash placed obliquely above and to the right. The flash placement serves to enhance in white colour the ligaments between the bases of the fin rays and the inter-clavicle junction (pectoral set) and the medial inferior puboischiadic projection and central ridge (pelvic set). The image on the right is a frame from a video taken using an FTIR (Frustrated Total Internal Reflection) system, in which the areas which are in contact with the surface are more radiant (whiter) in proportion to proximity to surface and pressure applied. In this case the fish was climbing a vertical surface under flowing water and was in a stationary phase of the process, the relatively high ambient light makes the FTIR enhancement low and so much of the natural colour of the fish is visible. The dark anterior region at the top of the fish in the right panel indicates this area is mostly held away from the surface, with for instance the posterior mouth in contact with the surface. The fish in the right panel was fluttering the posterior margin of its pectoral fins (which is a common behaviour – see main text). Flutter channels are visible which run from the anterior head area to the posterior of the pectoral fins outside the margin of the pectoral suckers and on the medial margin of the fin rays. Panels C and D: Two frames of a video of a Sewellia species fish climbing a vertical transparent acrylic surface. The white circles highlight areas which are stationary with respect to the fish movement. The orange lines outline the axial direction of the pelvic and pectoral suckers and associated features. The pelvic fin rays swing back and forth on each step of the gait and fan out like a grass skirt. The pectoral sucker unit swings at a lesser angle to the direction of movement (vertically upward in these figures).

## Discussion

The hill stream loach sucker system works in a similar way to a silicone rubber sucker in that it indefinitely maintains a hold to a surface under elastic tension with no energy input. Release and replacement involves the breaking, or partial breaking, of the seal and the equalisation of pressure between the inner space of the sucker and the outside environment. This is confirmed by the anatomic features earlier described leading to problems with preservation. Thus energy is expended in opening the sucker. The suction pressure exerted by the loaches in this study was around a tenth to a twentieth of the pressure exerted by an equivalent sized silicone rubber sucker. The loaches ‘walked’ up the vertical glass waterfall by pushing back with their fin rays in the same way as oars propel a rowing boat, and by sliding their suckers over the surface without breaking the seal. Here we falsify the hypothesis that these fish always use the suckers as the anchor points for movement (walking from sucker to sucker suggested by De Meyer and Geerinckx (2014)) and support the alternate hypothesis that they more often use their fins as the forward propulsion anchor and the suckers as constantly sliding surface attachment. Thus sliding friction of a sucker may be a determining factor for the maximum applied pressure. We further falsify the hypothesis that the suckers are used to overcome hydrodynamic lift in the climbing scenario (lift is theoretically a more likely scenario in moving water than downforce (Vogel 1993)). Our modelled dead specimen demonstrated that body shape alone would cause a slight downforce and maintain contact with the surface which supports the speculation of De Meyer and Geerinckx (2014). Thus our alternative explanation is that the mild suction pressure is used to counter the downforce required to use the fin rays as oars and maintain a positive pressure in the mouth region. We have shown that using this method a hill stream loach can overcome a drag equivalent to 3 to 4 times its own bodyweight while climbing a vertical glass waterfall.

The key control area for both sets of suckers is in the central ventral section; the suprapelvic flap opens and seals both the pectoral and pelvic sucker groups. Our results suggest that the pelvic fins are probably the primary anchor and propulsive unit during a vertical climb as they swing further and sustain higher pressures. The pectoral sucker group however is shown to be more skeletally complex, with more moving parts and with applied pressure correlated to size of fish. So the pectoral sucker is likely to exert more variable pressure and finer control for steering or reacting to a complex water flow field. Each of the 4 suckers can be opened and closed independently. The pectoral and pelvic pairs can be internally connected.

The resting periods reported here during climbing are made possible by the fact that the fish can continue to pass water through its buccal cavity and over its gills. We have shown that the fish does this by maintaining a constant positive pressure in the mouth area, and having this section entirely isolated from the suction pressure of the body sucker both externally and internally to its body. This is demonstrated here, 1) by observation of the of the CT scans showing the physical isolation of the breathing cavity by the elaborately adapted pectoral girdle, and 2) by the positive water pressure maintained under the anterior head area. The positive pressure around the mouth is not replicated with a passive model fish. These observations are supported by physiological analysis (De Meyer and Geerinckx 2014) and are further supported by the observation that the fish will climb damp surfaces without flowing water but were not observed to rest on dry surfaces.

The positive pressure under the head could be controlled and maintained by use of the flutter channels and associated fin fluttering. Fluttering is often observed while feeding and during other benthic activities, not associated with climbing (De Meyer and Geerinckx 2014), and the evidence of the way in which this system operates which is provided here leads to a set of new hypotheses that can be developed about function. This functional description extends and clarifies the suggestions made by De Meyer and Geerinckx (2014) about the way in which fluttering is used to distribute pressure and water under the body. Our evidence here suggests it does this by creating positive pressure under the head and in the suprapelvic flap area and that the flutter channels are used to maintain a particular level of positive pressure under the head isolated from the main suckers; this suggests the channels are used as pressure release valves independent of the mouth and suckers, and/or as pump inputs from the pectoral posterior margin fluttering.

Recent work on *Cryptotora thamicola* (the blind cave fish) suggested that it walked like early tetrapods and that it uniquely had a tetrapod-like pelvic girdle (Flammang et al. 2016). We mentioned in the introduction how the movements of the sucker loaches in general can be similar to the cave fish (a close relative and a loach), but can also be more diverse and complicated, including walking backward, upside down, and laterally. Firstly this work vindicates the approach taken by Flammang et al. (2016) to concentrate on the gait of the fins as the propulsive units as even the loaches with the most elaborately adapted suckers use their fins as the propulsive anchors. Secondly, there are some striking similarities in the skeletal structures of the loaches described here and the cave fish. However we have shown that the *Sewellia* sp. has a substantially different plate position and muscle attachment. The gait of the *Sewellia* sp. is often a diagonal-couplets lateral sequence gait like the cave fish (Flammang et al. 2016), although its muscular attachments and pelvic plate position are very different, and may be thus studied as a precursor to movement on land. But we suspect that the sucker loaches described here have a suite of possible gaits dependent on the wide range of movements described above and seen in the videos presented in the supplementary material (but also see Robert 1982 for descriptions of crawling as opposed to creeping). When the fish is climbing a near vertical wall under a torrent of moving water, our results show that the pelvic sucker group moves more than the pectoral set, and the tail moves more than the head. The maximum amplitude of the standing wave on the fish’s spine in this case is moved down the body and is probably aligned with the pelvic girdle. It seems likely that the standing wave gait of the sucker loach smoothly adjusts from crawling, through moving against shallow flow, to swimming in deeper water while remaining in ground effect contact with the substrate. It is interesting that the largest and most elaborately adapted representatives (*Sewellia* sp., *Gastromyzon lepidogaster*, and *Sinogastromyzon puliensis*) are physically endemic to areas very distant from each other but are all very similar in shape, size and habitat use, although (as we have shown here) have different skeletal structure. The compartmentalisation of the body into different pressure volumes by the skeletal development is an interesting adaptation, which is surprisingly closely mirrored in convergent evolution by many species in this family that have separately radiated across Southern Asia (Sawada 1982). Climbing ability may be the primary evolutionary constraint that has moulded these developments.

This work is also interesting because it has application to many objects that need to be dynamically attached to surfaces in flow. For example; tags to fish, temporary medical units in bodily vessels, and inspection devices on ships or aeroplane hulls. The propulsive unit has to overcome both sliding friction and gravity. The fish in this study push forward with their crescent shaped fin rays that fan out like oars and balance the suction pressure of their suckers with the pressure required to keep the fin rays anchored. The fish maintains overall contact with the substrate through downforce invoked by its body shape independently of this balance of forces required for locomotion. The convergent evolution of these morphologies across multiple radiations suggests it is uniquely well adapted to the engineering challenges presented to both animals and machines of this size in these conditions.

## Acknowledgements

Funded by the Leverhulme Trust.

## Supplementary material

Locomotion:

video 1 – sucker, adult Gastromyzon sparring
video 2 – ground effect, adult Sewellia sparring
video 3 – open water swimming
Photo 1 – Inverted movement, clinging etc.
Video 4 – Juvenile sparring
Analysis
Video 5 – video analysis GUI in operation
Figure: tracked point movement in body zones
SDL: 3D volumes, pectoral set, pelvic set.
CT Scan location: Freely available at http://doi.org/10.17639/nott.7009 (Accession numbers to follow after confirmation w.r.t. to publishing)

## Notes

http://doi.org/10.17639/nott.7009

